# Poison exon annotations improve the yield of clinically relevant variants in genomic diagnostic testing

**DOI:** 10.1101/2023.01.12.523654

**Authors:** Stephanie A Felker, James MJ Lawlor, Susan M Hiatt, Michelle L Thompson, Donald R Latner, Candice R Finnila, Kevin M Bowling, Zachary T Bonnstetter, Katherine E Bonini, Nicole R Kelly, Whitley V Kelley, Anna CE Hurst, Melissa A Kelly, Ghunwa Nakouzi, Laura G Hendon, E Martina Bebin, Eimear E Kenny, Gregory M Cooper

## Abstract

**Purpose:** Neurodevelopmental disorders (NDDs) often result from rare genetic variation, but genomic testing yield for NDDs remains around 50%, suggesting some clinically relevant rare variants may be missed by standard analyses. Here we analyze “poison exons” (PEs) which, while often absent from standard gene annotations, are alternative exons whose inclusion results in a premature termination codon. Variants that alter PE inclusion can lead to loss-of-function and may be highly penetrant contributors to disease.

**Methods:** We curated published RNA-seq data from developing mouse cortex to define 1,937 PE regions conserved between humans and mice and potentially relevant to NDDs. We then analyzed variants found by genome sequencing in multiple NDD cohorts.

**Results:** Across 2,999 probands, we found six clinically relevant variants in PE regions that were previously overlooked. Five of these variants are in genes that are part of the sodium voltage-gated channel alpha subunit family (*SCN1A, SCN2A*, and *SCN8A*), associated with epilepsies. One variant is in *SNRPB*, associated with Cerebrocostomandibular Syndrome. These variants have moderate to high computational impact assessments, are absent from population variant databases, and were observed in probands with features consistent with those reported for the associated gene.

**Conclusion:** With only a minimal increase in variant analysis burden (most probands had zero or one candidate PE variants in a known NDD gene, with an average of 0.77 per proband), annotation of PEs can improve diagnostic yield for NDDs and likely other congenital conditions.

## INTRODUCTION

Neurodevelopmental disorders (NDDs) affect 1-2% of children and display a wide phenotypic range that includes seizures, developmental delay, autism, and intellectual disability^1^. NDDs often result from highly penetrant genetic variation, and 1,493 genes are confidently associated with developmental disorders^2^. Exome and genome sequencing can lead to molecular diagnoses in NDDs, as they allow for detection of rare variants across many genes. In turn, a molecular diagnosis, particularly early in life, can provide many benefits to patients and their families^3,4^. However, despite rapid improvements in genomic testing, such tests frequently fail to reveal any clinically relevant variants; across most studies, the diagnostic yield of genomic testing for NDDs is at or below 50%^5^.

The vast majority of variants found by diagnostic testing are in coding exons that are part of standard gene annotations. Non-coding intronic variants are often ignored due to lack of functional annotation. However, analyses of transcriptome data have revealed functionality in otherwise non-annotated intronic regions^6^. One such category of functional sequences includes “poison exons” (PEs). PEs are alternatively spliced exons that result in the creation of a premature termination codon (PTC), either by direct inclusion or by frameshifting the transcript. The inclusion of PTCs cause translation to stall, thereby triggering nonsense-mediated decay (NMD) of the transcript and reducing protein levels in the cell. Some PEs are spliced into transcripts at relatively high levels during mammalian brain development in early stages and are hypothesized to regulate protein levels of the genes in which they reside^7,8,9,10^. Crucially, since PE usage leads to NMD, and therefore reduction of protein, genetic variation that alters PE inclusion can lead to a loss-of-function effect. Variant-induced PE inclusion is known to cause a small proportion of NDDs including *SCN1A*-associated epilepsies and Cerebrocostomandibular Syndrome (MIM 117650)^11,12^.

Despite their biological and disease relevance, PEs are often absent from standard gene models and PE-associated variants may be dismissed as irrelevant deep intronic variants. To remedy this, there have been reannotation efforts resulting in the addition of some of these regions to gene models. For example, epilepsy-related PE transcripts were included in Human GENCODE release 33^13^. However, causal PE variants are likely still missed due to lack of routine implementation in the variant analysis process. This may, in part, reflect the concerns with analyst and clinician “alert fatigue”^14^ due to the dilution of variant analysis with too many benign intronic variants.

We used published RNA-seq data from developing mouse cortices sampled from embryonic day 14.5 to 2 years old^15^ to define a list of evolutionarily conserved PEs that are differentially expressed in the developing brain for application in variant analysis pipelines. We filtered these results to highlight PEs potentially relevant to NDD, and within these curated regions found PE variants suspected to contribute to NDD in six probands from a cohort of 2,999. These variants were missed in conventional genomic analyses. While these six probands only represent 0.2% of the total probands analyzed, there is minimal additional effort required to analyze PE variants, which compares favorably to the increase in the yield of clinically relevant variants. Further, beyond the potential diagnostic benefit to the individual probands, discovery of these alleles is of more general translational value, particularly in light of drug-discovery efforts targeted to PEs.

## MATERIALS AND METHODS

### Curation of Clinically Relevant Poison Exons

To identify potential PE regions of interest, we used publicly available results of deep mRNA sequencing of mouse cortexes during development at E14.5, E16.5, P0, P4, P7, P15, P30, 4 month, and 2-year timepoints^15^. Yan and colleagues (2015) reported 77,949 elements alternatively spliced in the mouse cortex over developmental time. To curate these elements for conserved PE in humans each element was filtered for the following criteria (Figure 1):

**Figure 1.**
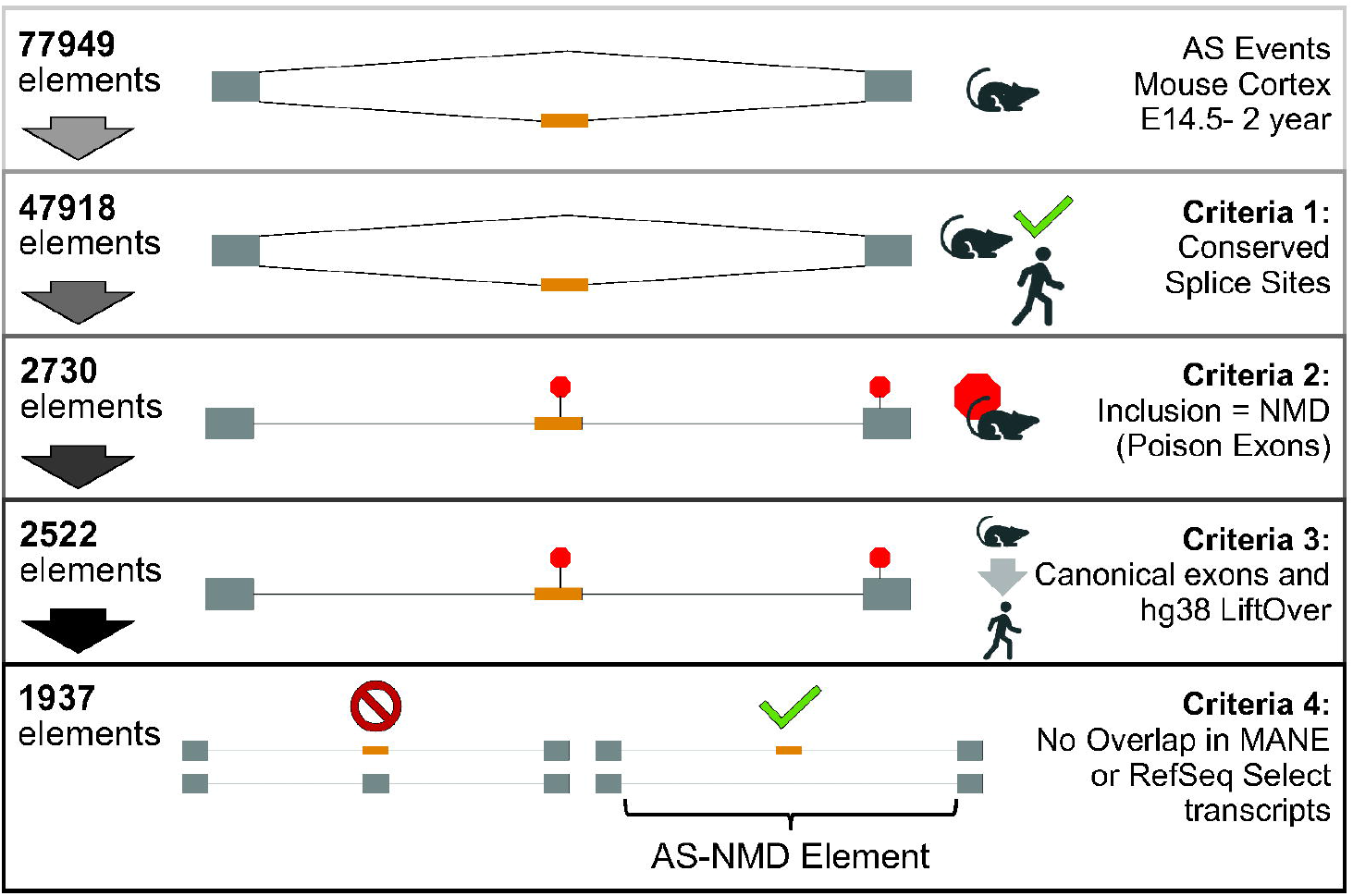
Extracting Clinically Relevant Poison Exons from Conserved Cassette Exons in Mouse Cortex. Graphic depiction of the methodology implemented to filter differentially spliced cassette exons in mouse cortex to find poison exons relevant to human neurodevelopmental disease. Poison exons are depicted in orange, and canonical exons are depicted in gray.

1. The AG/GU (+/- 1 or 2 canonical splice sites) are conserved between mm10 and hg19 (the reference assemblies used by Yan et al.,)
2. Alternatively-spliced (AS) element is determined to cause NMD upon inclusion (“NMD_in”) per Yan et al.
3. The cassette must be included between two canonical exons in mm10, and its coordinates must successfully lift over from mm10 into hg38
4. The hypothetical PE must not overlap with a Matched Annotation from the NCBI and EMBL-EBI (MANE) or RefSeq Select transcript exon

Data utilized for Criteria 1 and 2 were provided by the “Non_redundant_cass_mm10_summary.txt” supplementary file provided in the Yan publication at https://zhanglab.c2b2.columbia.edu/index.php/Cortex_AS. Criteria 3 was determined by intersecting the conserved NMD_in elements with the mm10 NCBI RefSeq UCSC track to select elements with distal 5’ and 3’ intron coordinates that overlap with the canonical exon scaffold^16,17^ using bedtools^18^. Criteria 4 evaluation was conducted by lifting PE cassettes into the hg38 genome using the UCSC Lift Genome Annotations tool^19^. These elements were then intersected with the NCBI RefSeq Select and MANE (nbciRefSeqSelect) dataset table from the UCSC Table Browser using bedtools. Each gene was assigned a disease consequence as determined by Online Mendelian in Man (OMIM)^20^. These elements and cassettes are available in Supplementary Data 1. Genomic coordinates of the PE cassettes are available in Browser Extensible Data (BED) format as Supplementary Data 2, and genomic coordinates of the introns containing the PEs are available in BED format as Supplementary Data 3. Supplementary Data 1-3 and the code and URLs to the external data used to curate these regions can be found at https://github.com/HudsonAlpha/poison_exon_variant_analysis.

### Cohort data acquisition

Data from probands in eight different research cohorts were analyzed, all part of translational genome sequencing studies on probands suspected to have congenital disease (Table 1). Data were acquired and results were returned in these investigations in accordance with site-specific informed consent and IRB-approved protocols. SouthSeq and NYCKidSeq are projects within the Clinical Sequencing Evidence-Generating Research (CSER2) Consortium^28^. Genomic data from SouthSeq, HudsonAlpha Clinical Sequencing Exploratory Research Consortium (CSER1), Alabama Genomic Health Initiative (AGHI), University of Alabama in Birmingham treatment-resistant epilepsy cannabidiol response clinical trial (CBD), Children’s of Alabama Genome Sequencing (COAGS), Alabama Pediatric Genomics Initiative (PGEN), and the University of Alabama in Birmingham Undiagnosed Disease Program (UDP)) were sequenced at HudsonAlpha Genomic Services Lab (GSL) or at the HudsonAlpha Clinical Services Lab (CSL). Genomic data from the NYCKidSeq cohort was acquired via the NHGRI Genomic Data Science Analysis, Visualization, and Informatics Lab-Space^29^ (AnVIL; see “NYCKidSeq Cohort Analysis on the NHGRI AnVIL”). Cohort-specific enrollment, sequencing, quality control, and analytic details are described in their respective publications (Table 1). Briefly, DNA was extracted from whole blood and sequencing libraries were constructed with or without PCR amplification. DNA library fragments were sequenced from both ends (paired) with a read length of 150 base pairs using the Illumina HiSeq or NovaSeq platforms to a target mean depth of 30x and target coverage of >80% of bases covered at 20x. Quality control included confirmation of each sample’s expected biological sex based on counts of chrX heterozygous variants and chrY variants and, when applicable, expected family relationships using somalier v0.2.5^30^ or KING v2.1.8^31^.

**Table 1.** Summary of Cohorts Screened. Probands were screened from previously published cohorts or clinical studies as indicated. All probands screened had suspected early-onset congenital NDD.

| Cohort Name | Reference | IRB Number | IRB Number |
| --- | --- | --- | --- |
| NYCKidSeq | 21,22 | Icahn School of Medicine at Mount Sinai IRB: (STUDY-17-00780/STUDY-20-01353);<br>Albert Einstein College of Medicine/Montefiore Medical Center IRB: (STUDY-17-00780/STUDY-20-01353) | 1032<sup>a</sup> |
| SouthSeq | 23 | University of Alabama in Birmingham IRB: IRB-300000328 | 636 |
| CSER1 | 24 | University of Washington IRB: 44776 | 532 |
| AGHI | 25 | University of Alabama in Birmingham IRB: IRB-170303004 | 484 |
| CBD | 26 | University of Alabama at Birmingham IRB: IRB-140905010/IRB-140826007 | 113 |
| Children's of Alabama | 27 | University of Alabama at Birmingham IRB: IRB 17-0314004 | 98<sup>b</sup> |
| PGEN | N/A (clinical study) | Western IRB: 00071 | 78 |
| UDP | N/A (clinical study) | University of Alabama at Birmingham IRB: IRB 170303004 | 26 |
| Total |  |  | 2999 |

<sup>a</sup>Total number of probands enrolled in study is 1052
<sup>b</sup>Total number of probands enrolled at time of writing is 146

For this analysis, all HudsonAlpha GSL/CSL samples were re-processed through a single sequence alignment and variant calling pipeline. Sequence reads were aligned to GRCh38.p12 using the Sentieon v201808.07 implementation^32^ of BWA-MEM^33^ and command line option -M -K 10000000. BWAKit was used for post-alt processing of the alternate contig alignments. Duplicate reads were marked, and base quality scores were recalibrated with Sentieon v201808.07 using dbsnp v.146 and Mills and 1000G gold standard indels as training data^32^. Variants were called on the hg38 primary contigs (chr1-chr22, chrX, chrY, chrM) using Strelka v2.9.10^34^ in germline single-sample analysis mode. For related samples, Illumina’s gvcfgenotyper v2019.02.26 (https://github.com/Illumina/gvcfgenotyper/) was used to merge the strelka genome VCF (gVCF) files into a multi-sample VCF to easily determine variant inheritance.

### NYCKidSeq Cohort Analysis on the NHGRI AnVIL

NYCKidSeq genome VCFs were collected and analyzed at one of two clinical laboratories (the New York Genome Center or Rady Children’s Institute for Genomic Medicine)^19,35^. Briefly, we collaborated with the CSER2 Data Coordination Center^36^ to develop a Docker image containing common genomics tools including bcftools v1.9^37^, htslib v1.9, and bedtools v2.30.0. We developed a workflow in Workflow Description Language (WDL) that uses bedtools and bcftools to quality-filter and region-filter VCFs equivalently to our local filtering process as described below in “Cohort Analysis with Clinically Relevant Regions.” The Dockerfile, WDL, and URLs for the deployed image and AnVIL workflow are available at https://github.com/HudsonAlpha/poison_exon_variant_analysis.

### Cohort Analysis with Clinically Relevant Regions

Variant call format (VCF) data from all cohorts was filtered for quality using the bcftools “filter” command requiring all of the following:

1. “PASS” variant calling filter status
2. Variant total read depth greater than 10 reads
3. Heterozygous genotypes having 20% to 80% of reads supporting the alternate allele
4. Genotype quality over 80

The resulting quality-filtered VCFs were region-filtered using bedtools intersect and a BED file containing the genomic coordinates of PE-containing introns detailed in Supplementary Data 1.

Annotations for the resulting quality-filtered and region-filtered variants were generated using the Ensembl VEP v102^38^. VEP plugins and custom annotations were used to add Combined Annotation Dependent Depletion scores v1.6 (CADD score)^39^, Genomic Evolutionary Rate Profiling scores (GERP score)^38^ scores, and variant frequencies from gnomAD v3.1.1^41^ and TOPMed Freeze 8^42^. Finally, variants were filtered for biological relevance via a custom R script to require all the following:

1. Variant consequence includes a non-coding effect (“NMD_transcript_variant”, “intron_variant”, “non_coding_transcript_variant”, or “non_coding_transcript_exon_variant”)
2. Variant is predicted in the top 10% of deleterious variants by CADD (CADD score ≥ 10)
3. Variant position is predicted to be evolutionarily conserved by GERP (GERP score ≥ 0)
4. Variant is absent from, or appears at most with an allele count of 1, in the TOPMed and gnomAD databases

Filtered variants were curated based on proband phenotype and disease relevance and variants of interest were classified using American College of Medical Genetics and Genomics (ACMG) and Association of Molecular Pathology (AMP) guidelines^43,44,45^. Variant annotation settings are available at https://github.com/HudsonAlpha/poison_exon_variant_analysis. Poison exon variants are reported in Table 2 with GRCh38 coordinates and HGVS nomenclature, validated in VariantValidator^46^.

**Table 2.**
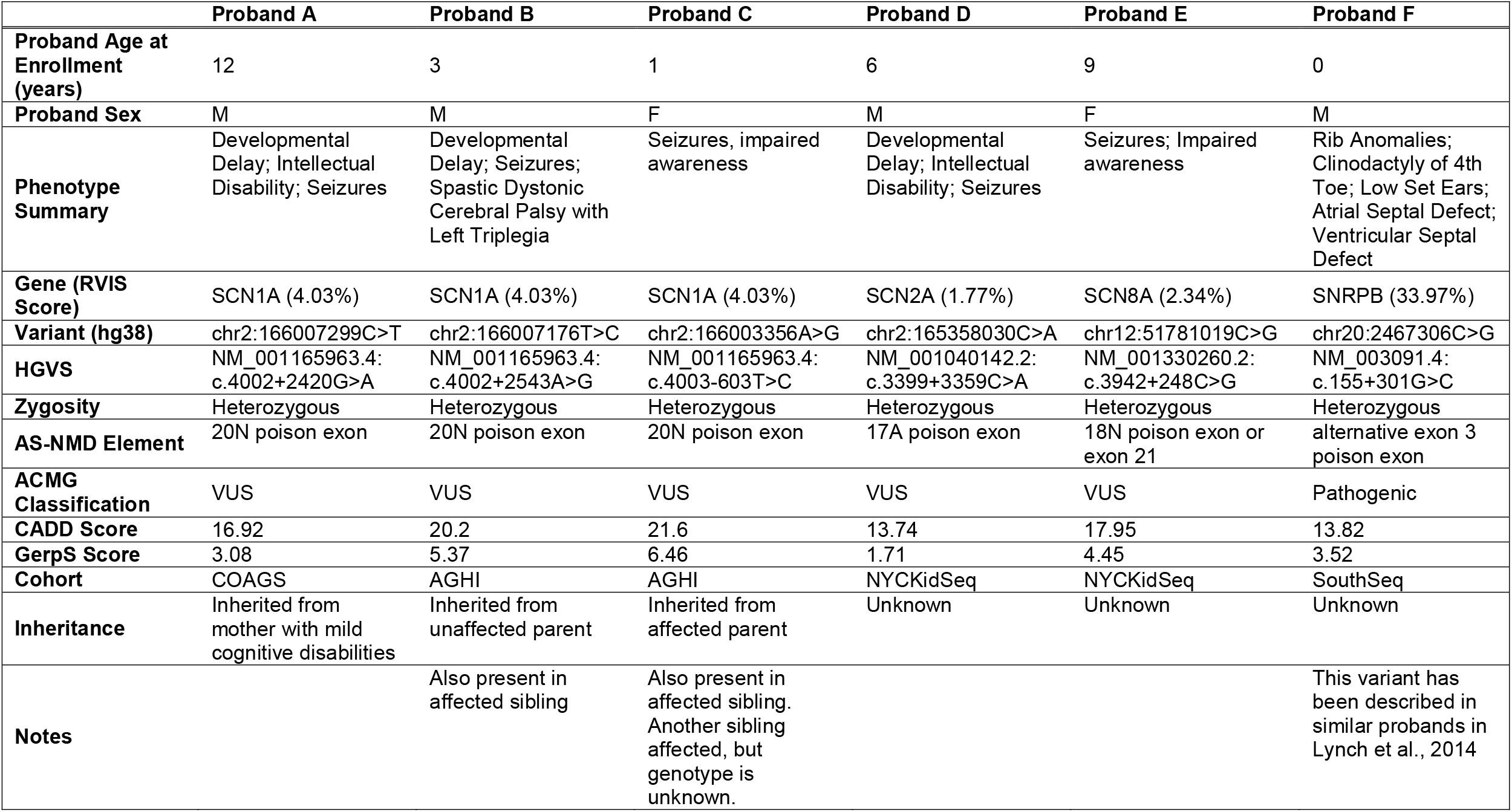
Poison Exon Variants Discovered. Five VUS and 1 Pathogenic variant were discovered in this analysis. CADD and GERP scores are shown for each variant, as well as the RVIS score for each gene in which the variant resides, along with select clinical and inheritance information.

## RESULTS

### Curation of Regions for Clinical Analysis

Yan and colleagues found 77,949 differentially spliced cassette exons in mice that resulted in alternative coding and NMD isoforms^15^. From these alternatively spliced exons, 47,918 had AG/GU splice sites conserved with human (hg19), and 2,730 of those resulted in NMD upon inclusion. 2,522 of these elements were included between two canonical exons in mm10 and successfully lifted over to hg38. PEs with overlap in the hg38 NCBI and EMBL-EBI MANE or RefSeq Select transcripts were removed to exclude human canonical exons resulting in a total of 1,937 candidate PEs (Supplemental Data 1). In this set, 571 are in genes that have one or more associated OMIM phenotypes. A BED file of introns and PEs defined for this analysis are available in Supplemental Data 2 and 3, respectively.

### Poison Exon Variants

The regions described above were used to find PE variants in the eight cohorts described in Table 1. Individual proband VCFs were filtered to extract high-quality, rare, conserved, predicted-deleterious variants that lie within PE elements and are not annotated to disrupt a coding exon. Manual variant curation resulted in identification of six variants with potential relevance to proband phenotype (Table 2, Figure 2). Using ACMG guidelines, five were classified as Variants of Uncertain Significance and one as Pathogenic. These variants had moderate CADD scores (median 17.4, range 13.7-21.6), indicating they rank among the top 5% of most highly deleterious SNVs in the human reference genome; we note that, as all these variants are outside coding exons and PEs are not used in the CADD model^39^, these scores may underestimate their deleteriousness. These variants have moderate to high GERP scores (median 3.99, range 1.71-6.46^40^) and are within regions of significantly elevated mammalian sequence conservation (Figure 2), suggesting they affect positions that have been under selective constraint during mammalian evolution. All six variants are absent from gnomAD^41^ and TOPMed^42^.

**Figure 2.**
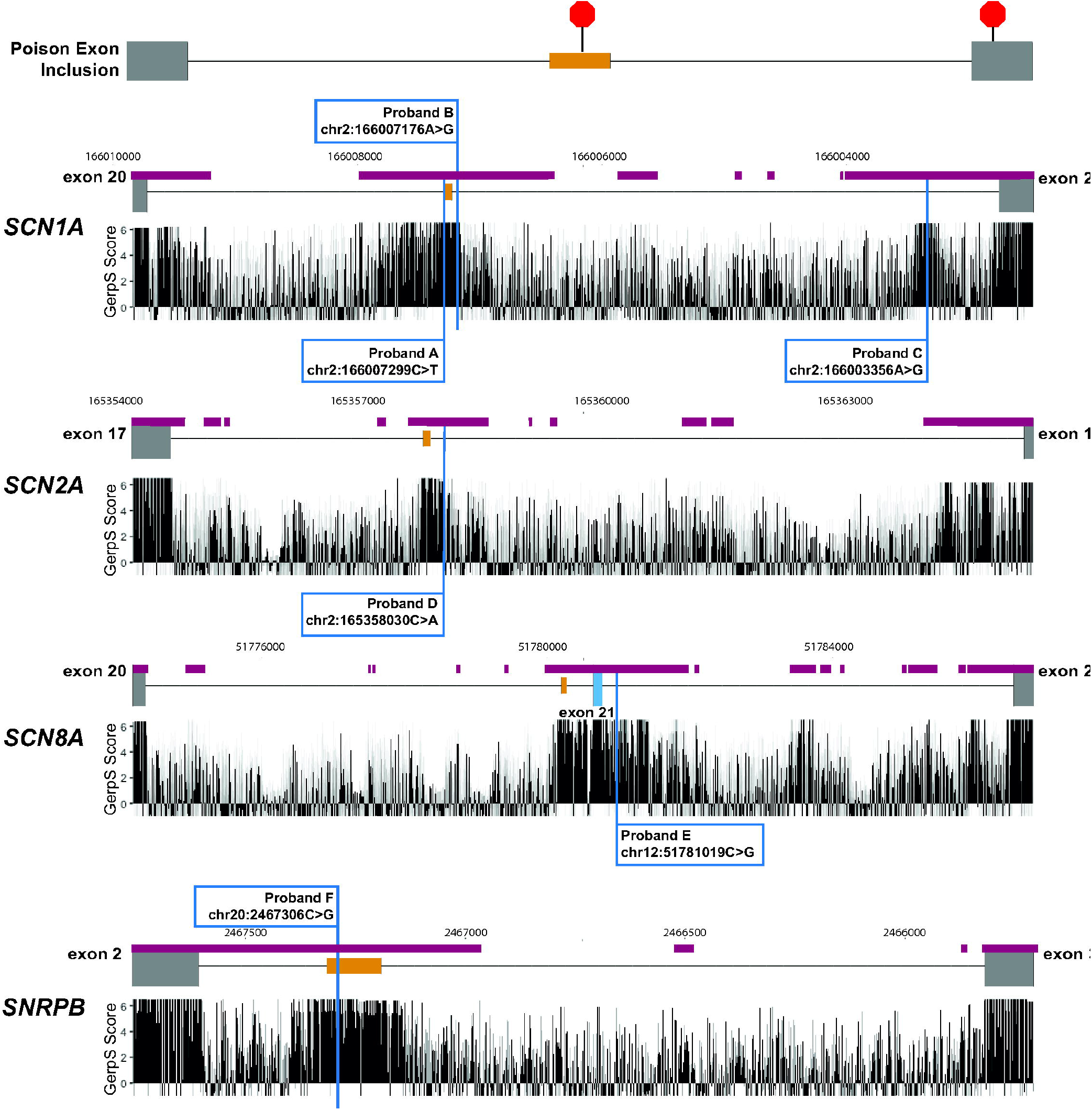
Variants found within introns containing poison exons in neurodevelopmental disease cohorts. Scale representations of the introns and canonical exons (gray), poison exons (orange), and alternatively spliced canonical exons (blue) in which variants were found. GERP scores are plotted below each and GERP conserved elements are noted (maroon bars).

Probands A, B, and C have variants within intron 20 of *SCN1A* and each has seizure phenotypes that are consistent with *SCN1A*-associated epilepsy (see Case Reports in Supplemental Data 4). This intron harbors the 20N poison exon^9^ variants in or near to 20N have been previously associated with epilepsy in both humans and mice^8,11^. The phenotype of Proband B also included developmental delay and spastic dystonic cerebral palsy with left triplegia. Proband C was reported to have intellectual disability and developmental delay. All three variants were inherited: Probands A and C inherited their variants from a parent affected with mild cognitive disabilities and seizures, respectively, and the variant found in proband B was inherited from an unaffected parent. Probands B and C had affected siblings that also shared their respective variants. Proband C has an additional deceased sibling who had seizures and whose variant status is unknown. The variants found in Probands A (chr2:166007299C>T) and B (chr2:166007176T>C) flank the 20N PE and are 7 and 54 base pairs away from the PE boundaries (chr2:166007230-166007293), respectively. The variant in Proband C (chr2:166003356A>G) is in a conserved region near the 5’ splice site of intron 20.

Proband D has a variant 170 base pairs from the 3’ boundary of the 17A PE (chr2:165358030C>A) of *SCN2A*. He exhibits developmental delay, intellectual disability, and seizures, which are consistent with *SCN2A*-associated phenotypes. Proband D had four VUSs reported from a targeted gene panel and a VUS reported from genome sequencing, but none of these variants were considered to be compelling in terms of disease relevance (see Case Report in Supplemental Data 4).

Proband E has a variant in a ∼1.5 kb conserved region containing both the 18N PE and the alternatively spliced exon 21 of *SCN8A*. Exon 21 is a canonical exon present in the MANE transcript of *SCN8A* but is skipped in 3 protein-coding transcripts (ENST00000668547.1, ENST00000545061.5, and ENST00000355133.7), and 18N is included in a truncated alternatively spliced isoform (ENST00000548086.3)^7^. She exhibits seizures and impaired awareness, consistent with Developmental and Epileptic Encephalopathy 13 (MIM 614558), associated with variation in *SCN8A*.

Proband F presented in the SouthSeq cohort with a phenotype consistent with Cerebrocostomandibular Syndrome (CCMS, MIM:182282), including rib anomalies, clinodactyly, ventricular and atrial septal defects, micrognathia, and facial dysmorphisms of deep-set eyes, low-set ears, and a prominent nasal bridge. The proband’s variant (chr20:2467306C>G) is within a PE in *SNRPB* and has been submitted to ClinVar as pathogenic (VCV000183431.3). This variant has been reported in 10 unrelated, affected individuals with both *de novo* and inherited alleles with incomplete penetrance^12^. Application of ACMG evidence codes results in a classification of Pathogenic.

## DISCUSSION

The diagnostic yield of genome sequencing for patients suspected to have congenital disease remains at or below 50%, and the incorporation of new genomic annotations may increase yield. However, additional annotations must be precise and biologically relevant so as to avoid diluting genome-wide analysis with an overwhelming number of inconsequential variants. Here, we curated a list of intronic regions of the human reference assembly that are: (1) associated with the PE-mediated regulation of gene expression in the developing mouse brain; (2) conserved between humans and mice; and (3) in genes associated with human neurodevelopmental and Mendelian diseases. We used these genomic regions to search for potentially disease-relevant variants that may affect PE inclusion in 2,999 probands from eight rare disease sequencing cohorts and discovered six variants of interest. These variants are all absent from the gnomAD and TOPMed variant frequency databases. The CADD scores for these variants are higher than 95-99% of human SNVs (range 13.74-21.6) but are at the low end of the range for variants associated with NDDs and other Mendelian diseases^47^. This may at least in part reflect the fact that these are deeply intronic variants in sparsely annotated regions and therefore lack most of the features used within the CADD model to infer deleteriousness. The primary annotation type supporting a deleterious effect is evolutionary sequence conservation. All these variants affect individual nucleotide positions with moderate to high degrees of mammalian conservation (GERP scores of 1.71-6.46) and are in regions of significantly elevated conservation^40^ (Figure 2).

Based on ACMG scoring criteria, five of the six variants we describe are VUSs, with only one evidence code applied based on absence from population databases (PM2). These five variants still require additional evidence to be assigned a more definitive status. Experimental analyses of transcripts showing evidence of altered PE inclusion are key, as this could provide experimental confirmation of a loss of function effect. Such work has been conducted for PE variants in *SCN1A* 20N^8,11^. In that context, we note that the three VUSs in *SCN1A* that we have identified here are all within the same intron containing the 20N PE studied by Carvill and colleagues (2018), who found five probands with rare variants in or near to 20N. Several of these variants led to altered 20N splicing *in vitro*^11^, and one was shown in mice to affect splicing of 20N *in vivo* and lead to phenotypes similar to those observed for *Scn1a* loss of function variants^8^.

A recent set of recommendations for interpretation of non-coding variants^44^ builds upon the previous recommendations^43,45^, but PEs are not specifically covered in this assessment. The authors suggested use of PS3_Strong in one example case where inclusion of a “cryptic exon” with a PTC was confirmed at the RNA level in *CFTR*. We believe PEs comprise a distinct subset of “cryptic exons” that warrant specific consideration. They are abundant across human genomes, including being present within hundreds of disease-associated genes. As the PEs described here were identified as alternatively spliced in mouse brain tissue, they comprise “biologically relevant” transcripts that are found in a key disease tissue of neurodevelopmental genes. Their high degrees of evolutionary sequence conservation further support their biological and disease relevance, and the potential value of defining PE-specific criteria for clinical variant assessment.

Given that loss of function is a well-established mechanism for many genetic conditions, and that by nature, PEs result in PTCs, this would suggest that application of the ACMG evidence code PVS1 may be appropriate. One previously reported variant near the *SCN1A* 20N PE, for example, was shown in heterozygous mice to lead to 50% reduction of SCN1A protein, as expected for a typical loss-of-function allele^8^. However, potentially weighing against usage of PVS1 is uncertainty in the magnitude of effect for a given PE variant and the relative dosage sensitivity of the affected genes. As noted by Ellingford et al., some PE variants may lead to only a partial effect wherein the variant allele produces both normal and truncated transcripts^44^. Further, the appropriateness of application of PP3 needs to be clarified for PE variants. For the five VUSs described here, we did not apply PP3 as the only available predictors included SpliceAI^48^ and CADD^39^, neither of which strongly support deleteriousness of these variants (SpliceAI maximum delta score of 0.00-0.31; CADD 13.74-21.6). This is likely due, at least in part, to the absence of PE annotations within the data used by variant impact prediction algorithms.

The pathogenic *SNRPB* variant in Proband F, while outside of standard coding exons, has been reported in at least 10 different patients, and has a confirmed loss-of-function consequence^12^. In the probands described here, it was initially filtered out as it has no coding effect annotation or other features that might protect it from removal. Its CADD score, for example, is only 13.82, which is lower than the vast majority of highly penetrant variants (and, as for other variants here, is likely deflated owing to the lack of features typically seen for deleterious variants). That said, a “rescue” filter that retains variants with P/LP designations in ClinVar, regardless of their coding status or other annotations, would lead to retention of this variant. Such a rescue would also, however, lead to additional variants in each proband requiring curation, with the magnitude of that addition dependent on the specific ClinVar parameters used (e.g., number and quality of submissions, presence/absence of conflicting interpretations, etc.).

Incorporating the 1,937 PEs and their surrounding introns detailed in Supplementary Data 1 adds minimal burden for variant analysis. These regions contained a median of four variants per proband (maximum of 37) passing through the criteria of having a CADD score of 10 or greater, a GERP score of zero or greater, and an allele count of zero or one in gnomAD or TOPMed. Among these, typically zero or one (median zero, maximum eight) variants were in a gene that has been confidently associated with a NDD phenotype via the Development Disorder Genotype - Phenotype Database^2^. For comparison, if one were to consider variants in all introns, a median of 94 variants per proband (maximum 1,057) would result from the same filtration, with a median of 17 variants (maximum 186) in genes associated with an NDD phenotype. To include all regions detailed in Yan et al., a median of 57 variants per proband (maximum 602) would result from the same variant filtration, with a median of 10 variants (maximum 100) in an NDD gene. Thus, our PE curation process reduces the analytical burden by 10-20X and yields variants with more evidence for disease relevance than more generic intron analyses, making their inclusion in clinical assessment practical.

The work described here provides a list of introns containing PEs that may be relevant to NDDs. Inclusion of these regions in genomic analysis resulted in identification of five novel VUSs; while uncertain, each of these variants is a compelling candidate for which functional follow-up may lead to reclassification. We also detected one Pathogenic variant that is likely to facilitate a clinical diagnosis. None of these six variants were previously detected and returned by previous genetic testing. The total increased yield among the 2,999 probands analyzed is thus ∼0.2%. Further, five of six of these variants were detected in probands with no returned variants, indicating the yield is mostly additive to current processes and is higher than 0.2% among probands with initially negative genome sequencing results.

Beyond increasing diagnostic yield, inclusion of PEs in analysis pipelines may prove beneficial to basic biological research and potential therapeutic research. More comprehensive discovery of patient-associated variation may facilitate improvements in computational effect predictions for variants in or near to PEs. It is also likely, given the abundance of PEs across the genome^15^, that new PE-disease associations remain to be discovered. Further, there are recent, ongoing efforts to use antisense oligonucleotide therapy (ASO) to target 20N inclusion in *Scn1a* via intracerebroventricular injection in mice at postnatal day 2^49^. These ASOs resulted in increased expression of *Scn1a* in brain tissue, and also a significant increase in survival of *Scn1a*-mutant mice. Phase 1 and 2 human clinical trials are being conducted for patients with Dravet syndrome (MIM 607208) using ASO therapy to target *SCN1A* mRNA containing the 20N poison exon (NCT04442295). A more comprehensive assessment of patients with PE variants is essential to understand the molecular and biological mechanisms of PE-associated disease, and thereby better understand the potential risks and benefits of PE-directed ASOs. Furthermore, it is possible if not probable that PE-directed ASOs may be more effective in patients whose disease results from a variant that affects PE inclusion compared to patients whose disease results from non-PE variation; identifying affected individuals with PE variants would be essential to build cohorts for clinical trials to assess this possibility.

In sum, our results provide substantial justification for including PEs in standard clinical genomic analysis to provide both short-term diagnostic and long-term research benefits.

## Supporting information

Legends for Supplemental Tables 1-3

## DATA AVAILABILITY

### Supplementary Data and Supplementary Methods

Poison exon data (Supplementary Data 1), hg38 PE cassette BED file (Supplementary Data 2), hg38 introns containing PE BED file (Supplementary Data 3), and variant extraction and supplementary methods including variant extraction, annotation, and filtering code can be found at: https://github.com/HudsonAlpha/poison_exon_variant_analysis

### Genome Sequencing Data

Genome sequencing and phenotype data are available for authorized access and hosted via dbGaP for the HudsonAlpha CSER cohort (phs001089.v4.p1). Genome and phenotype data will be available for authorized access via dbGaP and hosted on AnVIL for SouthSeq (phs002307.v1.p1) and NYCKidSeq (phs002337.v1.p1). Genome sequencing and phenotype data are not available for the AGHI, UDP, PGEN, UDP, or COAGS cohorts, as data sharing requirements were not incorporated into the funding mechanisms and/or consent processes of those cohorts.

## ACKNOWLEDGEMENTS

We thank all the families who participated in the studies detailed in Table 1. SAF is supported by NIMH F31MH126628. CSER was supported by grants from the US National Human Genome Research Institute (NHGRI; UM1HG007301). The SouthSeq project (U01HG007301) and the NYCKidSeq project (1U01HG0096108) were supported by the Clinical Sequencing Evidence-Generating Research (CSER2) consortium, which is funded by the National Human Genome Research Institute with co-funding from the National Institute on Minority Health and Health Disparities and the National Cancer Institute. AnVIL cloud compute credits were provided by grant support to the CSER2 Data Coordinating Center (U24HG007307). AGHI is conducted at the University of Alabama at Birmingham and the HudsonAlpha Institute for Biotechnology and funded by the state of Alabama. The Children’s of Alabama Board of Trustees funded the Children’s of Alabama Genome Sequencing (COAGS) study. The Alabama Pediatric Genomics Initiative (PGEN) cohort was funded by the Alabama Pediatric Genomics Initiative. We thank the University of Alabama at Birmingham Undiagnosed Diseases Program (UDP) and Dr. Bruce R. Korf. We thank the CSER2 Data Coordinating Center for their work facilitating data sharing and across the CSER Consortium and computation on the AnVIL, particularly Kathleen Ferar and Richard Green. We thank current and former employees of HudsonAlpha, HudsonAlpha Genomic Services Lab and the HudsonAlpha Clinical Services Lab who contributed to sequencing data acquisition and analysis, particularly J. Matthew Holt and David E. Gray.

## AUTHOR CONTRIBUTIONS

Conceptualization: S.A.F., G.M.C; Data curation: S.A.F., J.M.J.L.; Formal analysis: S.A.F.; Funding acquisition: S.A.F., E.E.K., G.M.C.; Investigation: S.A.F.; Methodology: S.A.F, J.M.J.L.; Project administration: S.A.F., C.R.F., G.M.C., Resources: E.E.K., G.M.C; Software: S.A.F., J.M.J.L., Z.T.B.; Supervision: G.M.C.; Validation: S.M.H., M.L.T., D.R.L., K.M.B., N.R.K., W.V.K., M.A.K., G.N.; Visualization: S.A.F.; Writing-original draft: S.A.F., J.M.J.L., L.G.H., K.E.B., E.M.B, A.C.E.H.; Writing-review & editing: S.A.F., J.M.J.L., S.M.H., G.M.C.

## ETHICS DECLARATION

All studies were conducted under the oversight of institutional review boards documented in Table 1. Informed consent was acquired from all probands and their families, and all information has been de-identified.

