## Supplementary material for "Poison exon annotations improve the yield of clinically relevant variants in genomic diagnostic testing": Legends for Supplemental Tables 1-3

**Supplementary Data 1-3 Legends**

**Supplementary Data 1. Annotated table of poison exon cassettes and their containing introns.** A excel spreadsheet with the following data:

1. PE_chromosome - genomic coordinates of the poison exon cassette
2. PE_start - genomic coordinates of the poison exon cassette
3. PE_end - genomic coordinates of the poison exon cassette
4. PE_identifier - unique identifier of the poison exon cassette. Format is PE_[sequential_number]_of_[total_number]_in_[intron_id] where intron_id pairs with the intron identifier in element_identifier column
5. Cassette_Yan_2015_Class - Classification of cassette from Yan et al. 2015
6. gene_symbol - HGNC gene symbol
7. alt_gene_list - list of alternative gene names
8. element_chromosome - genomic coordinates of the RefSeq intron containing the poison exon cassette
9. element_start - genomic coordinates of the RefSeq intron containing the poison exon cassette
10. element_end - genomic coordinates of the RefSeq intron containing the poison exon cassette
11. element_identifier - unique identifier of the RefSeq intron from the UCSC RefSeq RefSeqAll table
12. MIM_number - OMIM disease ID
13. MIM_phenotypes - OMIM phenotypes list
14. SFARI_genetic_category – SFARI Genetic Category
15. SFARI_gene_score – SFARI Gene Score
16. SFARI_syndromic – Syndromic Category (S or NA) if there exists independent evidence that the gene is associated with idiopathic ASD.

Columns 14-16 were sourced from the SFARI-Gene 20 July 2022 release accessed from https://gene.sfari.org/database/human-gene/ on 06 September 2022.

**Supplementary Data 2. BED file of poison exon cassette coordinates (hg38).** A BED file with the following data:

1. chrom: PE_chromosome - genomic coordinates of the poison exon cassette
2. chromStart: PE_start - genomic coordinates of the poison exon cassette
3. chromEnd: PE_end - genomic coordinates of the poison exon cassette
4. name: PE_identifier - unique identifier of the poison exon cassette. Format is PE_[sequential_number]_of_[total_number]_in_[intron_id] where intron_id pairs with the intron identifier in element_identifier column

**Supplementary Data 3. BED file of intronic regions containing one or more poison exons (hg38).** A bed file with the following data:

1. chrom: element_chromosome - genomic coordinates of the RefSeq intron containing the poison exon cassette
2. chromStart: element_start - genomic coordinates of the RefSeq intron containing the poison exon cassette
3. chromEnd: element_end - genomic coordinates of the RefSeq intron containing the poison exon cassette
4. name: element_identifier - unique identifier of the RefSeq intron from the UCSC RefSeq RefSeqAll table
