## Supplementary Data 4 for "Poison exon annotations improve the yield of clinically relevant variants in genomic diagnostic testing"

**Supplementary Data 4.** Detailed Case Reports of Probands with Poison Exon Variants Relevant to Proband Phenotype.

**Proband A**

Proband A is an 11-year-old male who presented to genetics at 8 years of age for developmental delay, borderline intellectual disability, and seizures. He was born at term, and he experienced gross motor delays and speech delays. He first had a febrile seizure at age 2 and had a total of four febrile seizures by age 6. At age 7 he had an additional febrile seizure and an abnormal EEG showed generalized spike and wave patterns depicting a poorly characterized epileptogenicity with two widespread bursts of irregular discharges less than 2 seconds in duration. He was then started on levetiracetam. At age 11, mother reports he began to have staring spells, but additional seizure activity during these episodes has not been confirmed. Prior head CT (age 6) and MRI (age 7) were normal. Neuropsychological testing at age 8 indicated concerns with attention, executive functioning, expressive and receptive language, and academic skills; he met diagnostic criteria for dyslexia. At age 10 he was diagnosed with autism, ADHD, and language impairment. Physical exam showed short upslanting palpebral fissures, slightly low-set ears, a long philtrum, and some mild joint hypermobility, but he is not overtly dysmorphic. Prior testing included normal microarray, Fragile X, plasma amino acids, and an acylcarnitine profile. At time of initial genome sequencing, the only reported finding was a maternally inherited variant of uncertain significance in *KDM5C* (c.3088C>T, p.Arg1030Trp); later methylation studies (EpiSign) were normal, indicating this was not consistent with a diagnosis. Genome reanalysis detailed in this investigation revealed the *SCN1A* poison exon variant, chr2:166007299C>T. His mother, who carries the same *SCN1A* 20N poison exon variant, reportedly had staring spells as a child; she struggled with reading in school and did not enter high school but has no formal diagnosis. A maternal uncle also has developmental delays but has not been evaluated by genetics. He has a sister with Prader-Willi syndrome due to a 15q11.2 methylation defect (normal microarray).

**Proband B**

Proband B is a four-year-old male who was first evaluated by genetics at age 2 for history of porencephaly, spastic dystonic cerebral palsy, and developmental delay. He was born at term with no neonatal complications. He experienced mild gross motor delays and was diagnosed with a receptive and expressive language delay. At the age of 2, he experienced a language regression where he stopped using words and now communicates by whining and pointing. He was evaluated by rehabilitative medicine at 10 months of age for left-sided weakness, and a brain MRI showed evidence of a right porencephalic cyst with associated ex vacu changes with HiNe global score of 49.5/78. He was diagnosed with spastic dystonic Cerebral Palsy with left triplegia. He was started on baclofen for tone and Botox. He had his first seizure at 2 years and 3 months of age, which involved right eye deviation, full-body stiffening, and lack of response to stimulation for 20 minutes. A repeat MRI redemonstrated the right porencephaly and showed loss of white matter, and he was started on levetiracetam. At age 2 years and 10 months he had another episode of eye deviation without a clear convulsive phase but had a period of altered awareness, and pyridoxine was added. Then at age 3 he was hospitalized for breakthrough seizures and oxcarbazepine was added to his regimen. At age 4, more breakthrough seizures occurred; he was switched from levetiracetam to clobazam, continued oxcarbazepine, and stopped taking pyridoxine, but 5 months later he experienced increased seizure frequency. He also has a history of myopia. Physical exam showed normal growth parameters, spasticity, and no dysmorphic features. Prior testing includes normal microarray, Fragile X testing, plasma amino acids, urine organic acids, acylcarnitine profile, and GAMT testing of blood and urine (testing for creatine deficiencies). He had genome sequencing at age 3 which was initially reported as negative, but genome re-analysis detailed in this investigation revealed the *SCN1A* 20N poison exon variant, chr2: 166007176T>C. This variant was also identified in his younger brother who has speech delay, autism, and history of febrile seizures (with a normal EEG at age 3), and their healthy mother who has no history of seizures or learning delays. However, their mother has a brother (maternal uncle to the proband, not tested for the presence of the variant) with a learning disability and possible autism.

**Proband C:**

Proband C is a 6-year-old female who presented at 8 months of age with a history of recurrent febrile seizures and developed partial seizures with impairment of consciousness that continued until the age of 4 years.  She has been seizure-free for 2 years on levetiracetam monotherapy, her brain MRI is normal, and EEG continues to show evidence of brief bursts of generalized spike and wave activity. Family history is significant for her father, who had seizures from 1 to 4 years of age which resolved, and her brother, who had a history of seizures and passed away at 23 months of age in his sleep, with cause of death at autopsy attributed to pneumonia. His age of initial seizure onset was 13 months, which was an episode of status epilepticus, and all his seizures were triggered by fever and/or illness. Proband C also has a younger sister with intractable partial seizures and history of febrile status epilepticus. Proband, father, and affected sister share the *SCN1A* 20N variant, chr2:166003356A>G, described in this study. Three paternal cousins have a history of febrile seizures, and one paternal cousin has generalized tonic-clonic seizures, but none of these relatives were available for genetic testing.

**Proband D:**

Proband D is a 6-year-old male with a history of epilepsy (prolonged seizures), intellectual disability, global developmental delay, and abnormal brain calcifications. His family history was unremarkable. Prior to enrollment in the NYCKidSeq study, the proband had two gene panels, both were uninformative. The proband received genome sequencing (GS) and a neurodevelopmental targeted gene panel (TGP) through the NYCKidSeq study. TGP identified four variants of uncertain significance, including a heterozygous, paternally inherited missense variant in *MAGEL2* (c.2498C>A, p.Ala833Glu, hg37). None of these were considered strong candidates for clinical relevance. GS also reported the paternally inherited *MAGEL2* VUS found via TGP but did not report the other VUSs found by the TGP. GS also identified heterozygous likely pathogenic/pathogenic variants in four genes associated with autosomal recessive disease: *KPTN, BTD, AP3B,* and *ATM.* While these findings did not change the proband’s medical management, reproductive risk was discussed with the family. It was also discussed that heterozygous pathogenic/likely pathogenic variants in *ATM* are associated with an increased risk for female breast cancer. GS was later amended to include sequencing of both proband’s parents. This amendment reported a heterozygous, *de novo* VUS in *CNNM2* (Chr10:102993371_102993372del, hg38). This variant is intronic and the observed phenotypes were considered a mismatch for *CNNM2*-associated disease (renal hypomagnesemia, MIM 607803). Of note, *SCN2A* was analyzed on the TGP but the chr2:165358030C>A poison exon variant resulting from this investigation was not reported, and inheritance of this variant is unknown.

**Proband E:**

Proband E is a 9-year-old female with a history of benign Rolandic epilepsy, with both focal and motor onset and impaired awareness. Her parents reported that at age four, she had four broken bones due to low calcium. She has identical twin brothers, age seven, with motor delays. Her paternal uncle, age 45, had a history of epilepsy as a child (approximately between ages 7-9) and has an arrythmia. The remaining family history was unremarkable. The proband received GS and a neurodevelopmental TGP through the NYCKidSeq study. TGP identified four uncertain variants which were not considered to be clinically suspicious by the research and clinical team and were therefore not returned. Trio sequencing via GS was negative. Of note, *SCN8A* was analyzed on the TGP, but the chr12:51781019C>G poison exon variant resulting from this investigation was not reported, and inheritance of this variant is unknown.

**Proband F**

Patient is a 4-year-old male who was born with mandibular hypoplasia, ten ribs with multiple posterior rib gaps, and an ASD/VSD.  Renal ultrasound was negative and head CT did not identify any intracranial abnormalities. Patient’s mother was counseled prenatally given abnormal ultrasound findings which included fetal micrognathia and a possible lower spine defect (meningocele). Genetic amniocentesis demonstrated a normal 46,XY fetal karyotype and negative microarray. Pediatric Genetics was concerned for cerebrocostomandibular syndrome (CCMS) after a postnatal exam which noted severe micro/retrognathia, deep-set eyes, low set and overfolded ears, high nasal bridge, and clinodactyly of the 4^th^ toe. Testing for *SNRPB* was negative on a multi-gene Craniofacial Panel performed by the Children’s Hospital of Philadelphia (CHOP). Patient has a healthy 6-year-old brother, and a maternal cousin deceased from complications of an open neural tube defect. The inheritance of the *SNRPB* poison exon variant resulting from this investigation, chr20:2467306C>G, is unknown, but is absent from his unaffected mother.
